# FInCH: FIJI plugin for automated and scalable whole-image analysis of protein expression and cell morphology

**DOI:** 10.1101/2024.04.20.590413

**Authors:** Cody A. Lee, Carmen Sánchez Moreno, Alexander V. Badyaev

## Abstract

Study of morphogenesis and its regulation requires analytical tools that enable simultaneous assessment of processes operating at cellular level, such as synthesis of transcription factors (TF), with their effects at the tissue scale. Most current studies conduct histological, cellular and immunochemical (IHC) analyses in separate steps, introducing inevitable biases in finding and alignment of areas of interest at vastly distinct scales of organization, as well as image distortion associated with image repositioning or file modifications. These problems are particularly severe for longitudinal analyses of growing structures that change size and shape. Here we introduce a python-based application for automated and complete whole-slide measurement of expression of multiple TFs and associated cellular morphology. The plugin collects data at customizable scale from the cell-level to the entire structure, records each data point with positional information, accounts for ontogenetic transformation of structures and variation in slide positioning with scalable grid, and includes a customizable file manager that outputs collected data in association with full details of image classification (e.g., ontogenetic stage, population, IHC assay). We demonstrate the utility and accuracy of this application by automated measurement of morphology and associated expression of eight TFs for more than six million cells recorded with full positional information in beak tissues across 12 developmental stages and 25 study populations of a wild passerine bird. Our script is freely available as an open-source Fiji plugin and can be applied to IHC slides from any imaging platforms and transcriptional factors.

**Graphical abstract:** 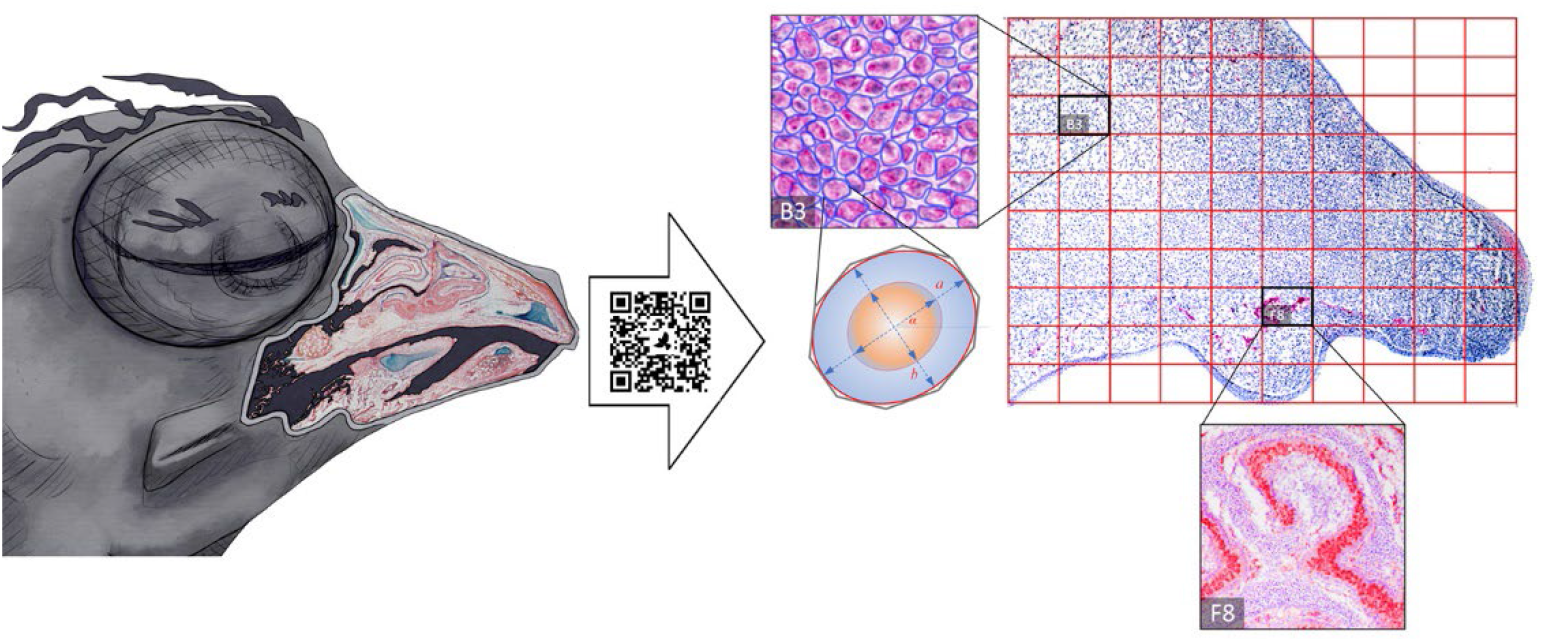

**Specifications table:** 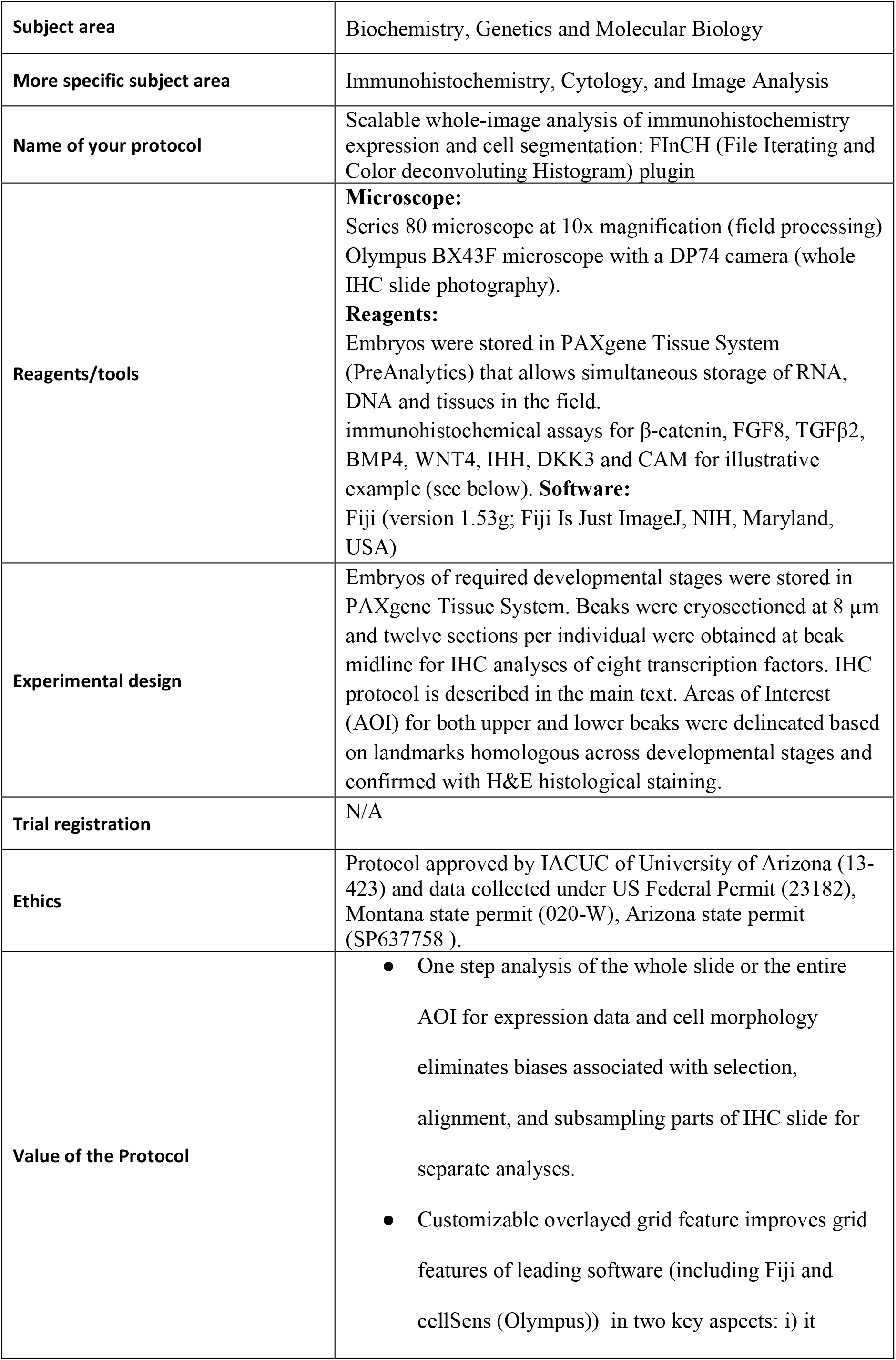

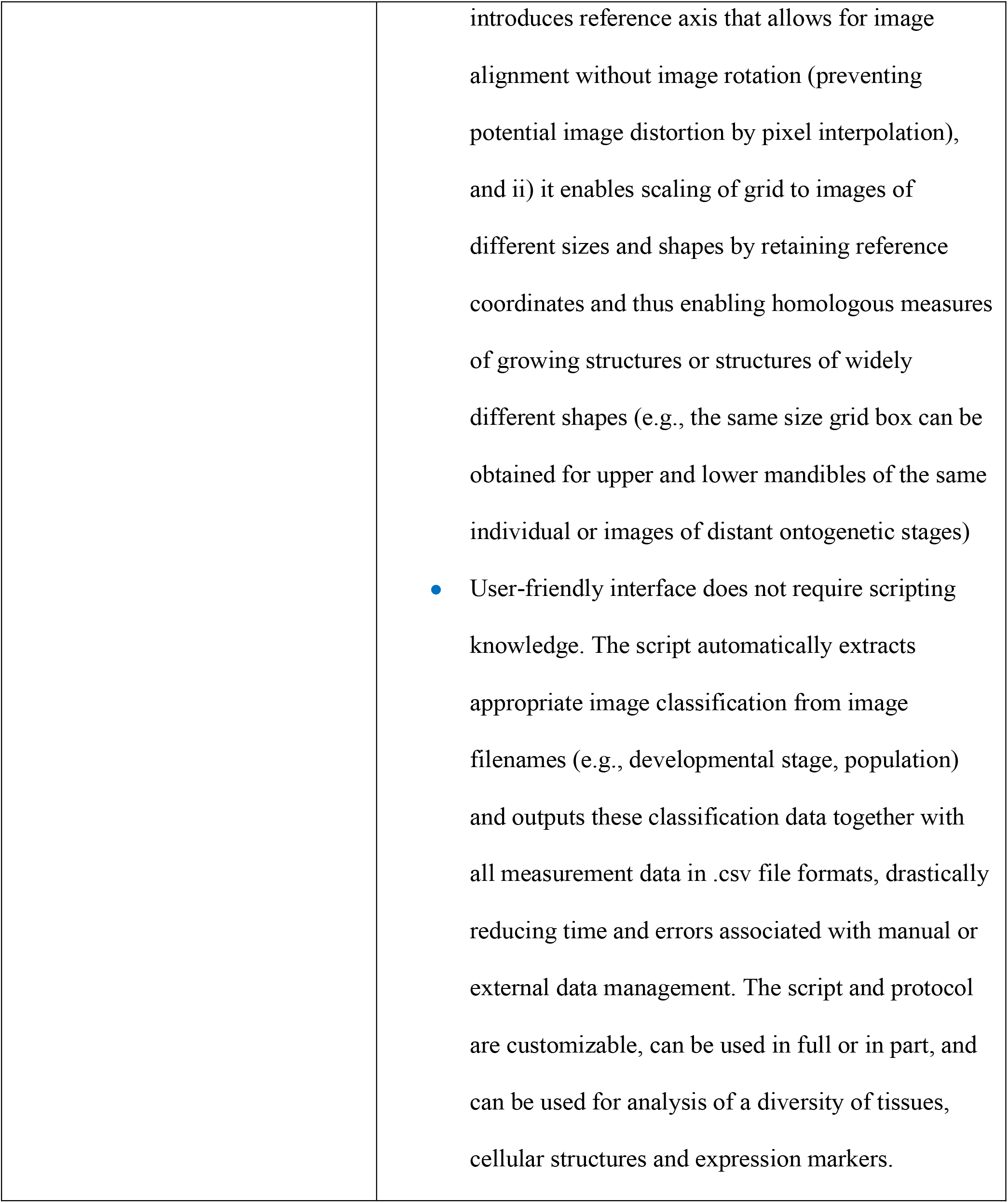

## Background

Evolutionary changes in ontogeny arise from modifications of time, rate and location of molecular regulation of developmental processes [1]. Immunohistochemical techniques are routinely used to investigate differences in expression of growth factors between developmental stages, populations, and species [2]. However directly linking these expression data with their downstream effects (such as the resulting changes in cellular morphology) has been challenging. Most studies examine these patterns either at the level of cells, such as associating nucleus morphology with TF production or at the meso-scale of the whole structure, selecting representative regions or tissues. What is needed is a scalable, whole-slide analysis that can enable compatible and repeatable measures across entire growing structures both within and across individuals. Here we introduce the protocol for the use of our python-based FInCH plugin – an automated, reliable, accurate, and high-throughput program that enables scalable whole-image analysis of protein expression and cell morphology. FInCH is freely available to download from Github at https://github.com/Lee-Cody/FInCH/.

### Description of protocol

We illustrate the use of this plugin with the study of cytological, histological, and regulatory changes driving microevolutionary diversification in beak morphology of the house finch (*Haemorhous mexicanus*) across 12 continuous developmental stages (HH25-HH36) and 25 study populations [3-5]. Here we specifically focus on measuring the timing and location of expression of eight key transcription factors (β-CAT, FGF8, TGFβ-II, IHH, WNT4, BMP4, DKK3, and CAM) [6-9] and associated changes in cell morphology, proliferation, and aggregation. This study system is particularly well-suited to highlight the advantages of our protocol and plugin for four main reasons. First, given recent evolutionary divergence of our study populations [10], we needed to accurately assess minute differences in time and location of regulatory expression and cellular modification across IHC images of full ontogenetic sequences that encompass significant changes in beak size and shape. Thus, the grid feature of our script would need to be scalable, and it should be possible to use the change in the grid scale itself to assess overall changes in beak size, in addition to identifying the same region of the structure and outputting exact location data for each cellular measurement and expression datapoint. Second, we needed to retain relative units of measurements for widely distinct structures within an individual but reset them between individuals. In our dataset, upper and lower avian beaks differ in shape and size, such that any area-specific measure (such as cell count or cell density or distribution of TF expression) must be retained within an individual (e.g., using column and row dimensions of upper beaks for lower beaks within each individual). Third, the method should allow precise and repeatable measurements for images that vary in horizontal alignment during initial imaging or slide mounting. Identical alignment is difficult to accomplish for growing structures with changing reference points. At the same time, the rotation of digital images during analyses introduces minor distortions caused by pixel interpolation, especially at the cell-level of measurements. Such distortions accumulate with repeated realignments at separate steps of TFs or cell and tissue measurements. Our script produces grid referencing without rotation or manipulation of original images and accommodates variable angles of imperfect horizontal alignments. Finally, the size and complexity of our dataset (>350 animals x eight TF accessed across 12 ontogenetic stages and 25 populations) required a high-throughput tool to analyze thousands of whole-slide images and millions of cells with high repeatability and fully indexed output. Our previous analysis of this dataset, utilizing manual subsampling of whole images for different analyses and grid features of commercially available software produced a large dataset [3], but was still limited to 450 x 10^3^ cells. Using FInCH, with the protocol described here, we now obtained full expression and morphology data for > 6 x 10^6^ cells in this dataset while requiring a fraction of the time and direct supervision.

### Experimental procedures

Our protocol utilizes high-resolution IHC expression images of the Area of Interest (AOI). Our illustrative sample are tissue sections of PAX-embedded beaks that underwent IHC for β-CAT, FGF8, TGFβ-II, IHH, WNT4, BMP4, DKK3, and CAM proteins throughout their early ontogeny (from stage HH25 to HH36). However, the protocol below is not specific to these proteins or structures. Briefly, embryos were stored in PAXgene Tissue System (PreAnalytics) and beaks were cryosectioned at 8 μm, and twelve sections per individual were obtained at beak midline for IHC analyses. Sections were blocked with an endogenous peroxidase (2% H_2_O_2_) for 20 minutes, washed in tris-buffered saline (TBS), blocked with 2% bovine serum albumin (BSA) plus 5% normal goat serum (NGS) in TBS for 1 hour. Avidin-Biotin blocking kit (Abcam) was used per manufacturer’s instructions before applying antibodies. Samples were incubated for 18 hrs with a primary antibody at 4°C, washed in TBS, incubated at room temperature for 1 hour with a secondary antibody, and washed again with TBS. Appropriate dilution of the primary antibody was determined by running preliminary trials with serial dilutions. IHC protocol is given in Supplementary Note 1. IHC sections were photographed at 600 dpi resolution with a DP74 camera using an Olympus BX43F microscope under 4x magnification and saved as tif files. File naming convention is important for correct extraction of data classification and automation of file operations and is described below. From each image, AOIs were extracted for the upper and lower beaks using H&E histological staining and homologous landmarks, saved as a separate tif file, and placed in AOI folder maintaining file naming convention (Fig. 1 shows workflow). A detailed description of AOI extraction is given in Supplementary Note 2.

**Fig. 1.**
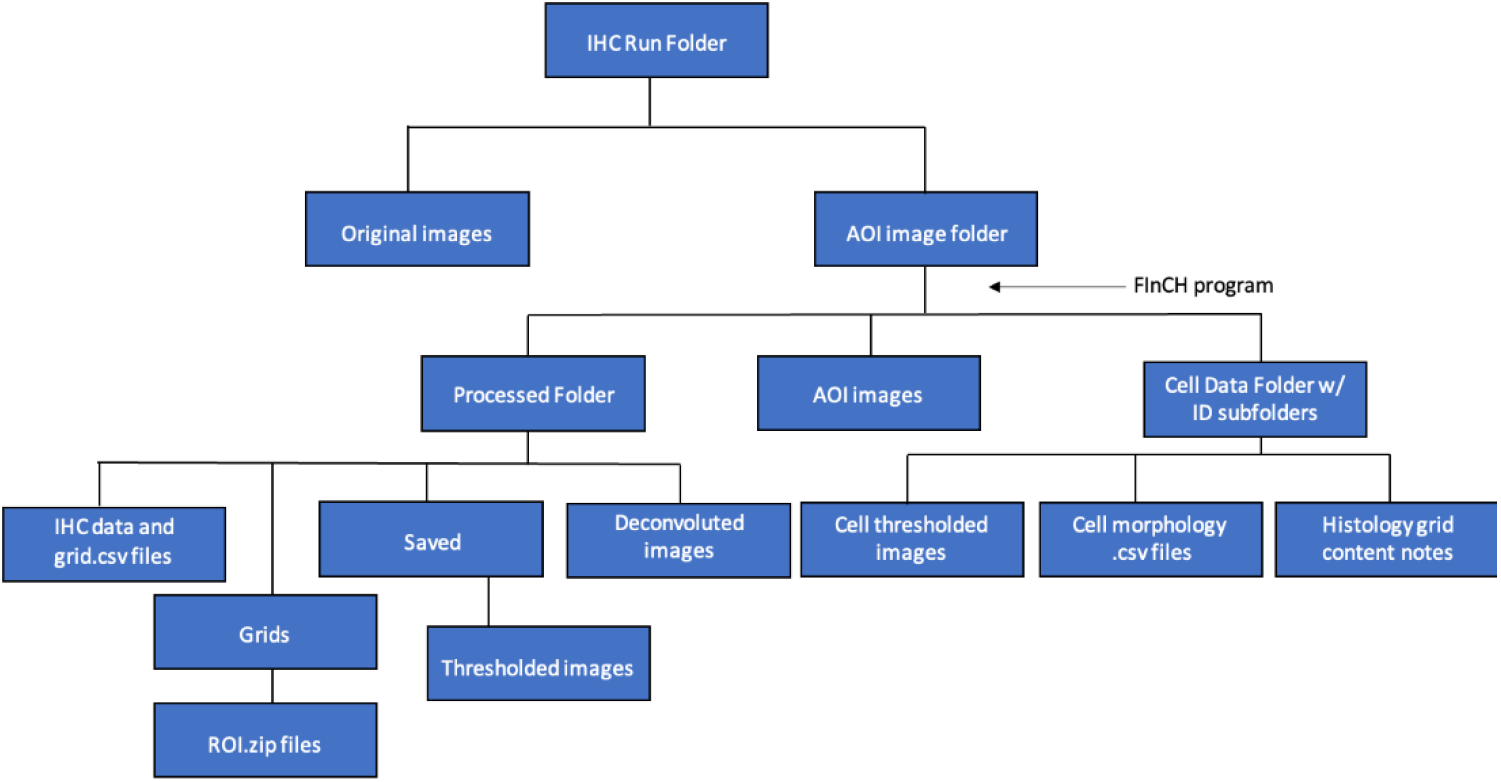
FInCH program workflow. The program processes the images in the “AOI image folder” (selected by the user). A “Processed folder” is created inside the “AOI image folder” and stores the deconvoluted images as well as the thresholded images (a folder “Saved”). The program also creates “IHC data and grid .csv files” folder in the “Processed folder”. “ROI.zip files” folder is created inside the “Grids” subfolder in the “Processed folder”. All cell morphology .csv files for each grid cell corresponding to the same image, are stored inside a folder with the image’s name inside the “Cell Data” folder. Finaly, the thresholded image that was used by the program to perform cell morphology analysis is saved along with the cell morphology.csv files inside the ID subfolder in the Cell Data folder.

### Synapsis of protocol

The plugin automatically and sequentially retrieves all AOI images from the folder, creates a customized scalable grid that detects image size with its orientation based on requested user input (and can retain the grid for repeated measures of the same image or homologous structures), includes a step for proofing of grid and alignment, retrieves IHC specification for TF from image name and applies appropriate color deconvolution, finds all cells expressing the specified TF, outputs converted TF expression images to a subfolder retaining names and TF, summarizes and outputs .csv expression data for each grid box (retaining spatial location) and overall structure, measures cell morphology for all cells identified in each of the grid boxes, and creates a subfolder for each image where it outputs .csv files with cell data for each grid box with location information. The plugin can be used for individual components of these analyses, for selected grid boxes or customized for other analyses requiring repeatable location-specific measurements. For example, the grid feature can be used for description of tissues and structures located within each grid box, associating TF and cell measures with histological conversion, specific anatomical structures, or tissue types.

### AOI Images

For optimal performance, images to be processed by FInCH should not contain structures outside of the immediate area of interest. The details of our AOI extraction are provided in Supplementary Note 2. Although not strictly necessary for FInCH to function, this step helps ensure accuracy of data collection and AOI border detection without close manual proofing. Any non-black or non-white pixels outside of the AOI on the image may be included in data collection or may interfere with accurate grid formation. One potential alternative may be to balance image colors via post-processing so that all background information is perfectly white (RGB: 255,255,255) or perfectly black (RGB: 0,0,0). This alternative method presents a risk of introducing inaccuracies within the AOI due to post-processing and has not been tested.

### Colour Deconvolution Spectra: separating stains on images

FInCH uses the built-in FIJI [11] module, Colour Deconvolution, to separate stain spectra on images [12]. The colour spectra are specified within a text file “colourdeconvolution.txt” inside FIJI’s plugins folder (Fiji.app/plugins/colourdeconvolution.txt). While FInCH can apply any of the default colour spectra options that come installed with FIJI, we highly recommend that users identify their own colour spectra for FInCH to use for colour deconvolution. This is because image properties vary by equipment and software, which can lead to inaccurate color separation if default settings are used.

Instructions for determining image spectra to be used for color deconvolution can be found at: https://blog.bham.ac.uk/intellimic/g-landini-software/colour-deconvolution-2/#newvectors. *Instructions for adding your own colour spectra* --. When adding your own spectra to the colourdeconvolution.txt file, make sure to follow the example shown at the top of the file (“#Stain_Name,R0,G0,B0,R1,G1,B1,R2,G2,B2”). Using the stain name, “user_stains1”, and the rgb values: (0.1111,0.2222,0.3333,0.4444,0.5555,0.6666,0.7777,0.8888,0.9999), add the following line anywhere in the colourdeconvolution.txt file (without the quotation marks) while making sure to leave the example line at the top of the file: “user_stains1,0.1111,0.2222,0.3333,0.4444,0.5555,0.6666,0.7777,0.8888,0.9999” When adding your own colour spectra, make sure that the RGB values for the stain you want analysed are used as the middle 3 values of the spectra colours (R1, G1, B1 in the example stain line). FInCH will keep, and process, the colour placed into the middle 3 values and discard the others.

### FInCH File Structure

FInCH has a modular design where functions and classes of similar themes are grouped into separate modules (Fig. 2). FInCH_.py is the manager script, DataManager.py is the core modulator with most of the programming functions.It performs initial setup, reads and creates settings as well as prior data files, processes images, and stores data. Interface.py manages most of the dialogs and menus. GridGen.py contains several FInCH classes with their associated methods. Utils.py includes supportive functions as well as the cell analysis function. FInCHPlugin.py is a placeholder that offers a blank space for users to add their own functions. It currently contains an example of how to extend FInCH with additional functions. At the initial start, FInCH will create a config file to track user settings (FInCH.ini). It also relies on the colour spectra file used by the FIJI module, ColourDeconvolution: “colourdeconvolution.txt”.

**Fig. 2.**
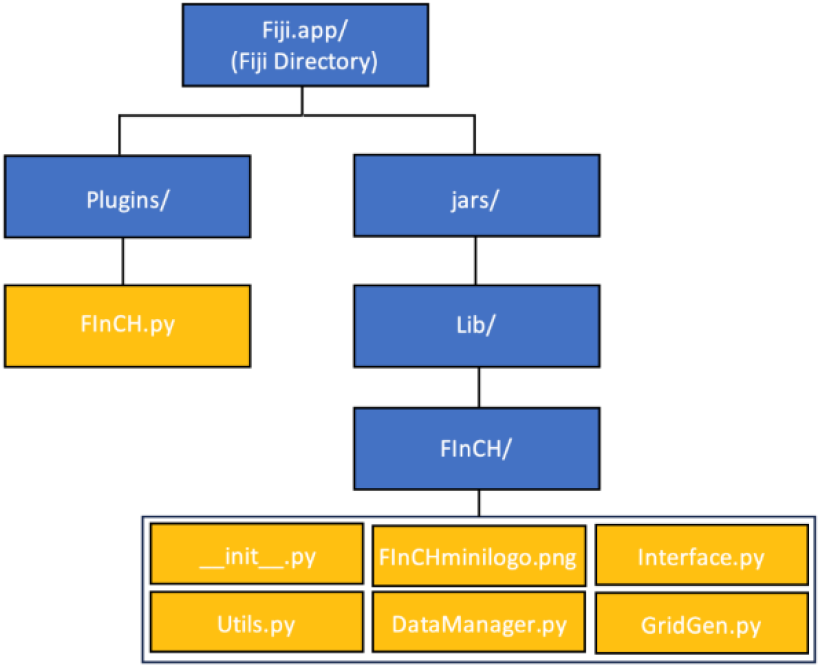
Structure of files included in FInCH plugin. Boxes in blue describe the folders that hold python scripts, boxes in yellow show associated scripts for FInCH Fiji functionality. FInCH_.py should be placed in the Fiji plugins folder (Fiji.app/plugins/). All the other files (init .py, Interface.py, DataManager.py, GridGen.py, Utils.py, FInCHPlugins.py, FInCHminilogo.png) should be placed in the jars/Lib folder (Fiji.app /jars/Lib/FInCH/).

### Initial Setup

When starting FInCH for the first time, it will present a dialog to select individualized user settings. The dialog will go through a few options that allow for users to change how FInCH identifies files, how it applies color spectra for color deconvolution, the threshold sensitivity of colour deconvolution, and customizable options, such as whether FInCH should look for specific designations (e.g., “upper” and “lower”) in filenames to preserve grid dimensions across multiple images (see below). Although there is also an option to use default settings to skip the setup dialogs, we recommend that users go through the setup dialogs at least once.

### Image naming

This program relies on the following image naming convention for data classification and integration. An underscore is used to separate elements in an image name.

**[Filename identifier]_[data classification]_[magnification]_AOI.tif**

1. [Filename identifier]: FInCH will use a list of text entries to identify images and to match these images to the correct colour deconvolution spectra (found in Fiji.app/plugins/colourdeconvolution.txt. Filenames should start with one of these identifiers followed by an underscore “_”. Filename identifiers are not case sensitive. In the protocol used here, FInCH has been used to analyze the expression of eight proteins: β-CAT, FGF8, TGFβ-II, IHH, WNT4, BMP4, DKK3, and CAM (Table 1). These are retained as examples in FInCH’s default settings and may be changed during the setup dialogs.
2. [Data classification]: Any classification data to be associated with data output, such as population, family, developmental stage, IHC run or slide, treatment). Separated by an underscore “_”.
3. [Magnification]: Filenames require specified magnification. In example protocol here images were captured under a 4x magnification, so the filename includes “4x”.
4. [_AOI.tif] ending concludes data classification string and tells the program that file is tif format.
5. Additional customization. In our example protocol, we analyzed paired structures for the same individual – upper and lower beaks. This options requests FInCH to retain grid coordinates and apply the same grid resolution (number of rows and columns) between associated structures of the same individual. This customization function must be enabled at the setup and “upper” or “lower” have to be included in the filename. FInCH will create a grid for the “upper” image first and then it will apply a grid with that resolution to the associate “lower” image. If a folder of “AOI” images includes files with “lower” in the filename, then an associated image must exist in the same folder with “upper” in the filename.

**Table 1.** File identifiers for transcription factors used in this protocol.

| Transcription Factor | Filename identifier |
| --- | --- |
| BMP4 (Bone Morphogenetic Protein 4) | Bmp4 |
| β-CAT (β-catenin) | BCat |
| FGF8 (Fibroblast growth factor 8) | Fgf8 |
| TGFβ-II (Transforming Growth Factor Beta Receptor 2) | TgfB2 |
| IHH (Indian Hedgehog Signaling Molecule) | Ihh |
| WNT4 (Wingless-related integration site 4) | Wnt4 |
| DKK3 (Dickkopf-3) | Dkk3 |
| CAM (calmodulin) | Cam |

Examples of file naming conventions with classification data to be associated with measurements in output data files:

**TgfB2_HH32_POP-1-N-A_s17_10_upper_4x_AOI.tif** (Analysis of **TGFβ-II** expression in an embryo at developmental stage **HH32** from a nest **1_N**, from egg **A** in population **POP**, IHC slide **s17**, run **10, upper** mandible, imaged under **x4**).

**Ihh_HH31-32_PLA-4-N-C_MID_s09_57_upper_20x_AOI.tif** (Analysis of **Ihh** expression in an embryo at developmental stage **HH31-32**, from a nest **4-N**, egg **C**, from **MID** section of **upper** jaw, IHC slide **s09**, run **57**, imaged under **20x**).

### Image analysis

#### 1. Collection of TF expression data and generating grid

1.1. Install FInCH_.py in the plugins folder of Fiji. [Fiji.app/plugins/FInCH_.py] (Fig. 2)
1.2. Install the “Libs” folder with all the FInCH modules (_init_.py, Interface.py, DataManager.py, GridGen.py, Utils.py, FInCHPlugins.py.) and FInCHminilogo.png in the “jars” folder of Fiji. [Fiji.app/jars/Lib/FInCH/] (Fig. 2)
1.3. If Fiji is running, close the program.
1.4. Open Fiji.
1.5. Start FInCH from the plugin’s menu. If it is the first time you open FInCH, an initial setup menu will appear (Fig. 3). The pop-up menu offers the option to use FInCH default settings which include the parameters shown in Fig. 4 (A-D) or change the initial setup by clicking on “Setup FInCH”.
1.6. Default settings (Fig. 5): Option E is selected; Option C is unselected. Click OK. FInCH will ask you to select the AOI folder. Non-AOI images can be inside this folder, but all AOI images should be in the AOI subfolder in this case (see notes). For optimal workflow and accuracy, all AOI images from the same IHC run should be in the same folder.
1.7. FInCH will then open all AOI images in that folder and will prompt user to draw a line on each AOI image. Press the TAB to cycle between open images and click OK when finished. This line is necessary to indicate the angle of the beak. The length of the line or its exact position is not important, but the angle should match the angle of the beak. Draw the line from the base of the beak to the tip. So, if the tip of the beak is on the lower right of the image and the base is on the upper left of the image, the line should be drawn from upper left to lower right.
1.8. After clicking OK, FInCH will process the images and save the data. When finished, it will open all processed images and the Image Navigator window (Fig. 6).
1.9. A new folder “Processed” will be created, containing all the colour deconvoluted images (Fig. 7b), as well as the thresholded images (Fig. 7c) stored in the “Saved” subfolder. Two .csv files will be generated and saved inside the “Processed” folder, data.csv, that contains the TF expression data, and grid.csv, with grid information that is required for reapplying grids to images or for repeatability assessment. It contains pixel coordinates for grids, matched to corresponding image titles. Along with these files, a subfolder named “Grids” is created, storing an ROI.zip file for each image, and used in cell morphology analysis.
1.10. Verify that the data.csv and grid.csv files (see file note explanations below) have been created for the entire run in the AOI/Processed folder. Select “Close All Windows”.

**Fig. 3.**
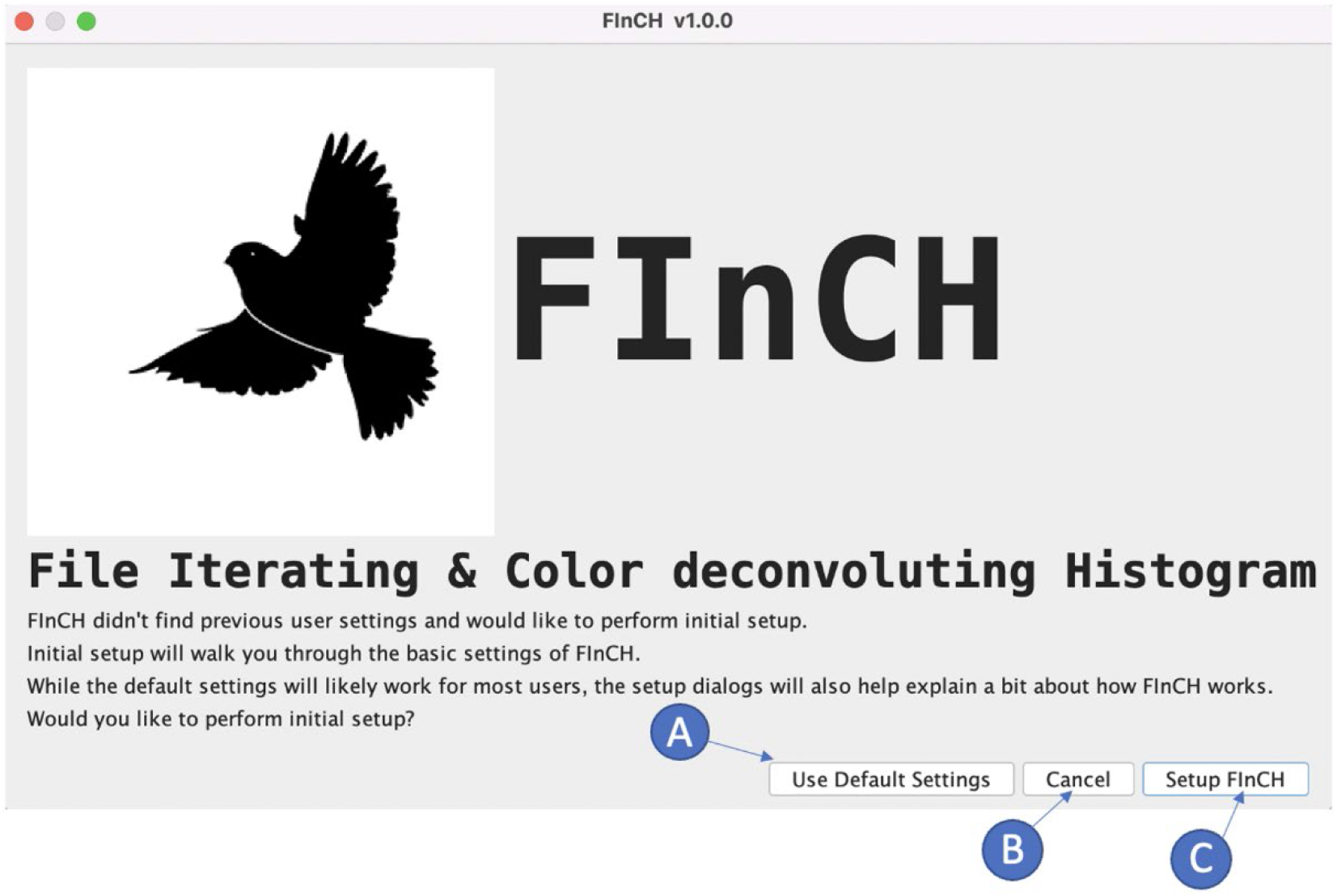
FInCH initial setup menu. If no additional changes are needed, select “Use Default Settings” (A). To make changes or additions select “Setup FInCH” (C). To terminate the plugin, select “Cancel” (B).

**Fig. 4.**
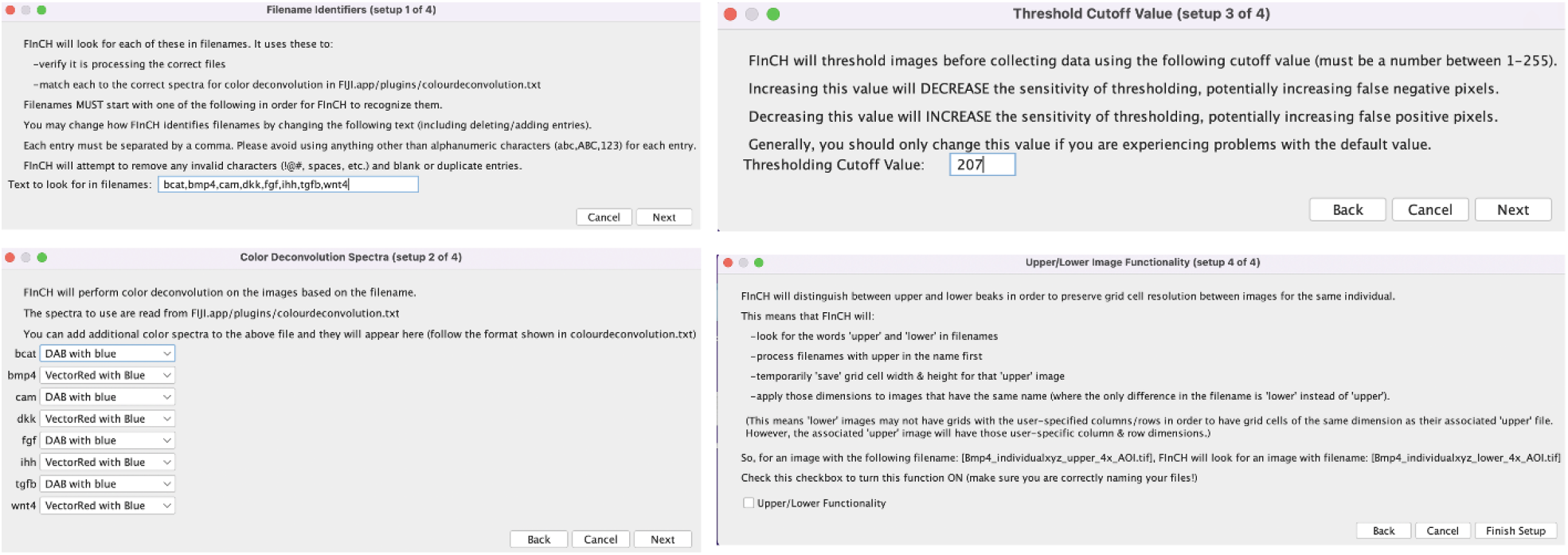
Setup parameters in FInCH. (A) Enter “Filename identifiers”, e.g., TFs for analyses. (B) Enter “Color Deconvolution Spectra” with the set of TFs analysed along with their IHC spectra as default. (C) Enter “Threshold Cutoff Value” (default value = 207). (D) Enter additional functionality, (e.g., “Upper/Lower Image Functionality”) or leave unchecked. This function creates a grid on a lower beak image with cell dimensions that match the grid cell of its associated upper beak image. Selecting “Finish Setup” to save new settings.

**Fig. 5.**
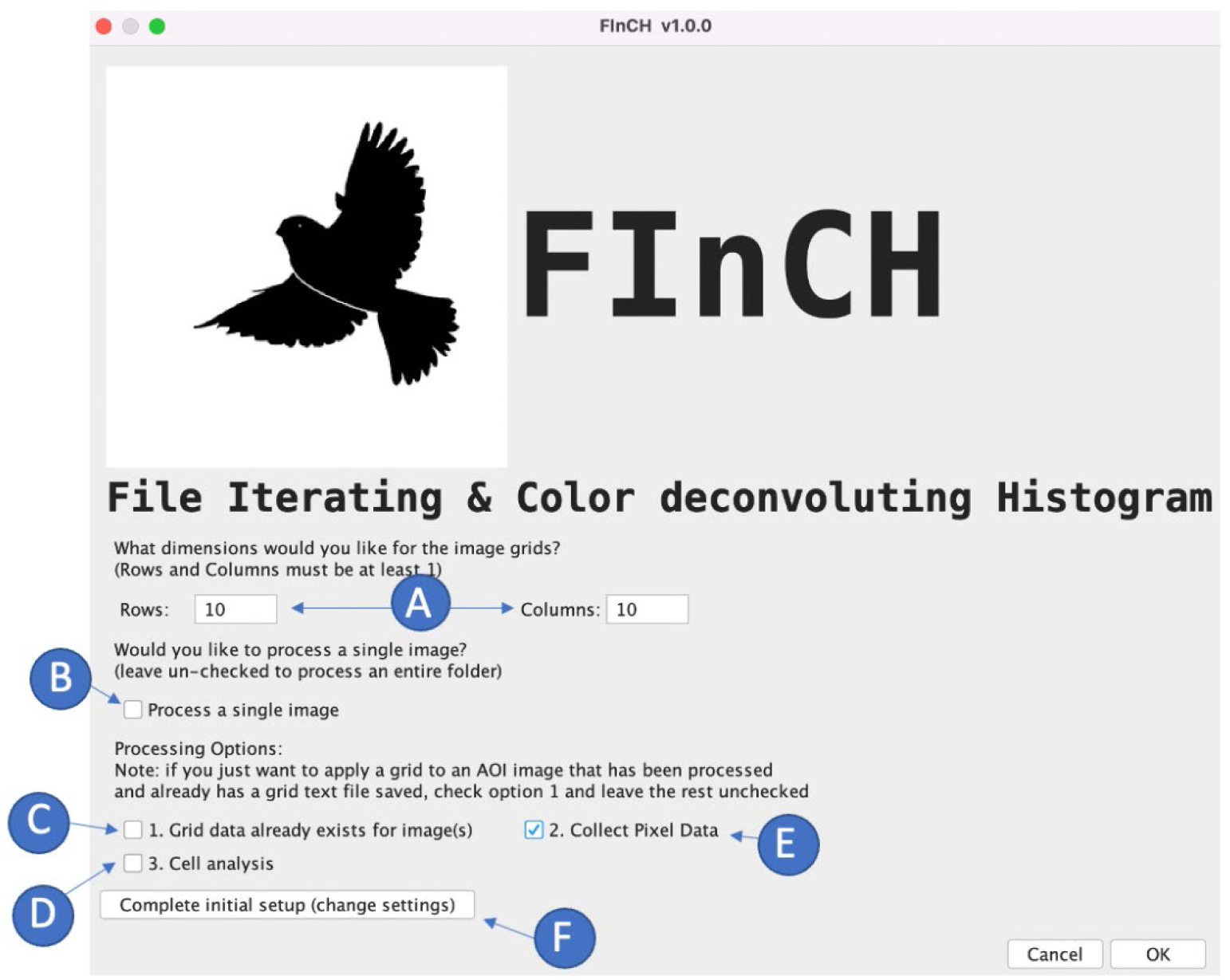
FInCH menu with the default settings. (A) Enter rows and columns to generate a grid (default 10×10). (B) “Process a single image” is unchecked by default (program assumes an entire folder will be processed). (C) “Grid data already exists for image(s)” is unchecked by default. If the user would like to open previously processed images with the previously created grid, this option should be selected, and the rest of the options should be left unchecked. The program also assumes that the user wants to do expression analysis (“Collect Pixel Data” (E) options checked by default). D-F Settings create a .zip file for each image, with the coordinates of the grid cells that will be necessary for future cell analysis. For cell morphology analysis, “Grid data already exists for image(s)” (C) and “Cell analysis” (D) options should be selected. (F) To modify FInCH settings, select “Complete initial setup (change settings)”

**Fig. 6.**
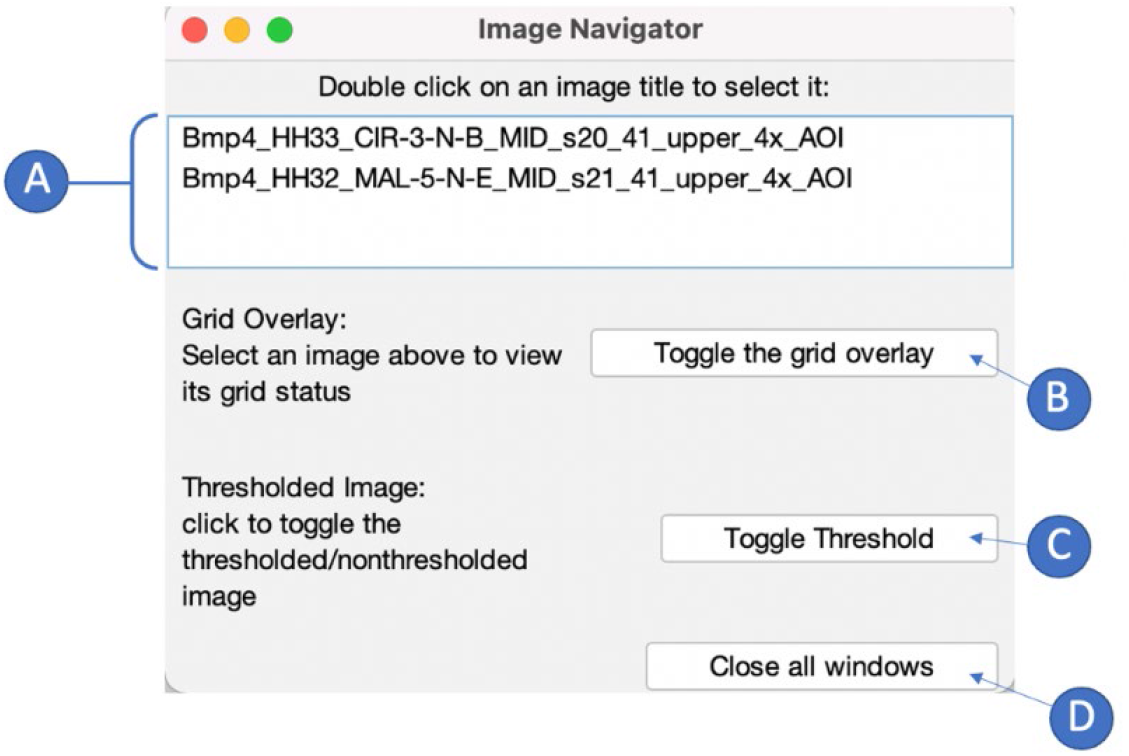
Image Navigator window once FInCH is finished processing images. (A) The names of the processed images are in the navigator window. Clicking on the filename will bring the corresponding image to the front of the screen. (B) “Toggle the grid overlay” or (C) “Toggle Threshold” will overlay the grid/threshold created by FinCH on the selected image. (D) “Close all windows” closes all windows opened by FinCH during the analyses.

**Fig. 7.**
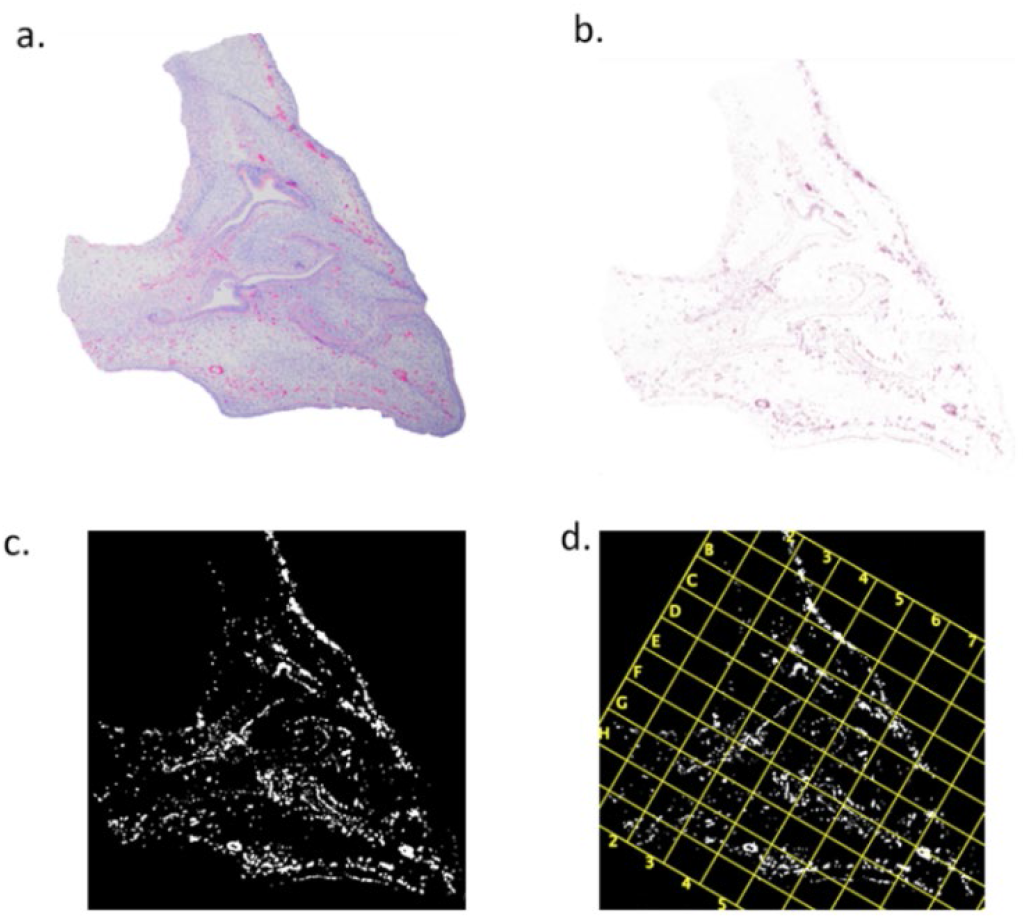
Representative stages of the TF expression analysis process in FInCH. (a) original image is (b) color deconvoluted by FInCH based on specified TF – only the TF expression is thresholded. (c) the thresholded image is analyzed by Fiji using the ROI Manager menu to compute the number of white pixels (which correspond to the TF expression) and black pixels (no TF expression) for (d) each grid cell. All stages of image processing can be manually confirmed by the user (Fig. 6).

#### 2. Reapplying grids to previously processed images

2.1 Start FInCH and check options shown in Figure 8. Click OK and FInCH will ask you to select the AOI folder that contains the AOI images to which you would like to reapply grids (e.g., the lower beak of the same individual when the grid was created for the upper beak).
2.2 FInCH will look for a grid.csv file with grid information for the images in the folder.
2.3 If a grid.csv file is found and the information in the file matches the image names in the AOI folder you selected, it will apply that grid information to the images and open all images. You can toggle grid visibility on each image using the Image Navigator window.
2.4 Reapplying the grid generated for the image allows for two additional tasks: i) histological or anatomical description of grid content (e.g., identification of structures or tissues) and ii) manual exclusion from data collection the grid boxes that contain damage due to sectioning or IHC.

**Fig. 8.**
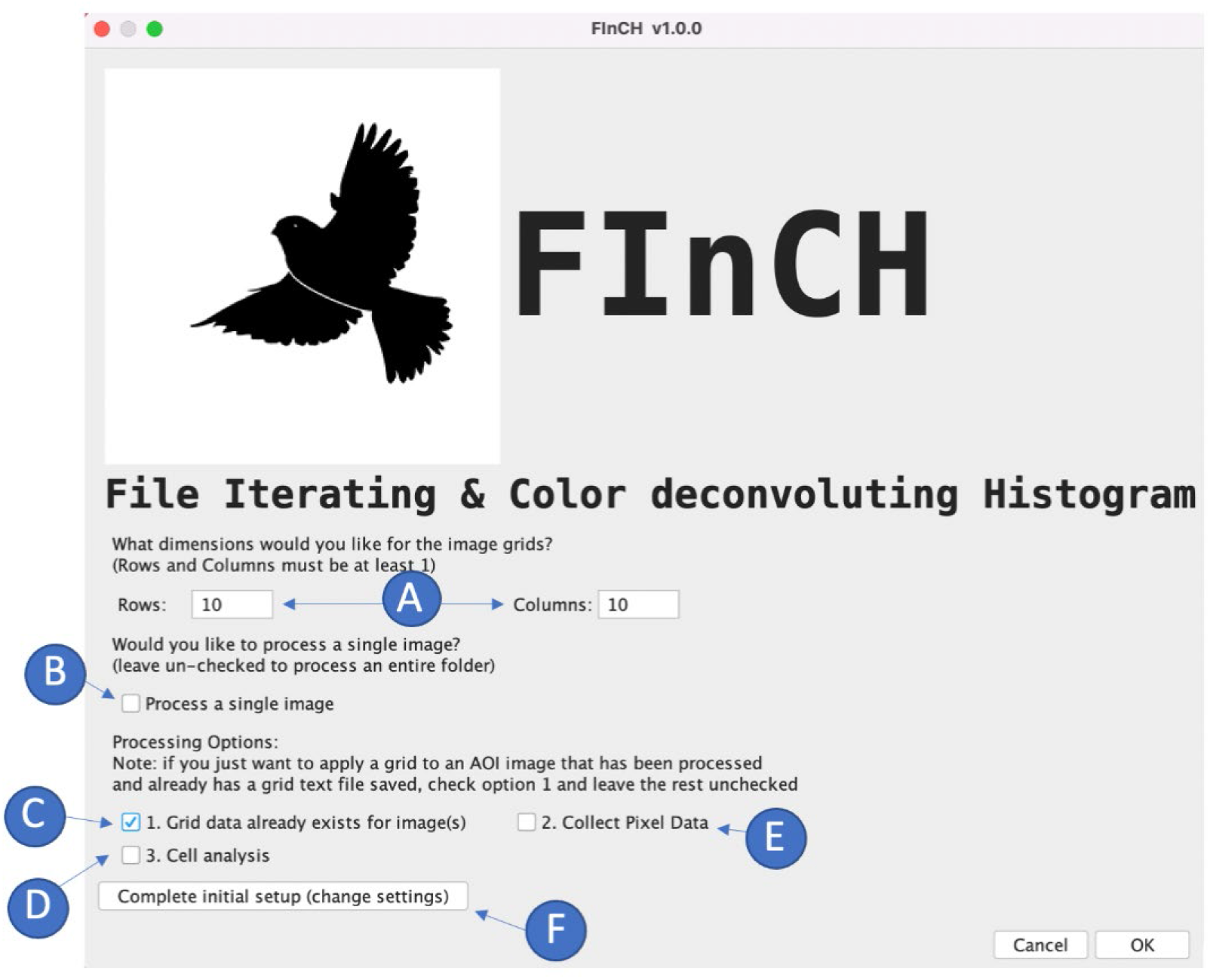
FInCH menu with settings for reapplying grid over processed images. All options should be unchecked except for option “C”, “Grid data already exists for image(s)”. Option “B”, “process a single image” can be both checked or unchecked, depending on if the user desires to process a single image or an entire folder respectively.

#### 3. Cell morphology analysis

3.1. Open Fiji and select the FInCH plugin in the plugins menu. Check the options shown in Figure 9.
3.2 Select the folder of AOI images for cell morphology analysis.
3.3 FInCH will now convert the images into 8-bit versions and apply custom script to subtract the image from its background, enhance boundaries, and apply an intensity threshold and watershed to define cell boundaries (Fig. 10).
3.4. Default Fiji scale or custom scale (1.293 pixels/micron in our sample) can be selected.
3.5. FInCH loads grid coordinate data from the previously created grid.csv file.
3.6. All cells in each grid box will be measured using the analyze particles feature of the FInCH program.
3.7. Any area of grid boxes that is outside of the image is not analyzed by the program, to ensure accurate data collection.
3.8. FInCH creates a “/cell_data” folder in the AOI image folder and subfolders corresponding to ID of grid box and retaining individual ID and all classifying markers included in image name (Fig. 1).

**Fig. 9.**
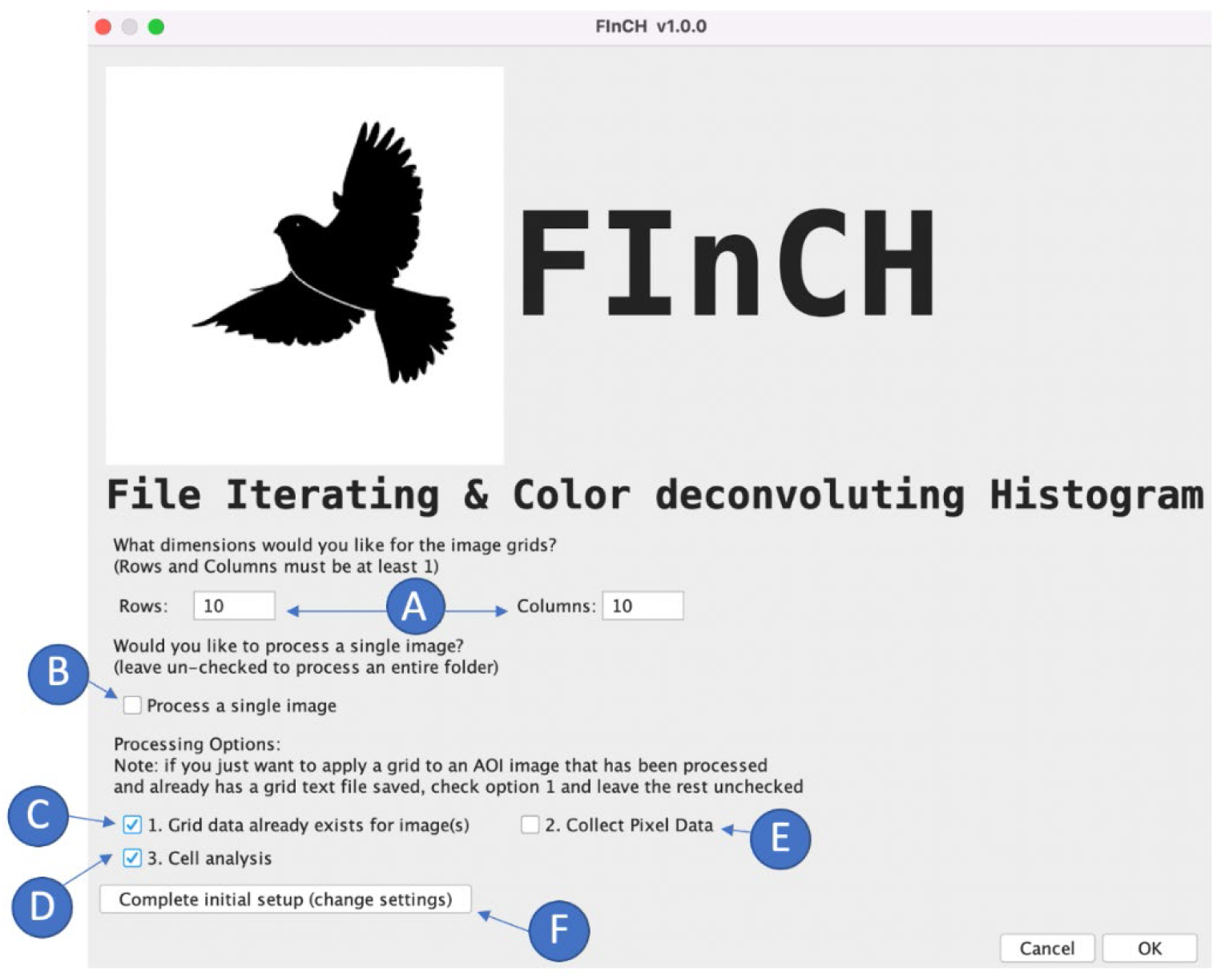
FInCH plugin menu for cell morphology analysis. All options should be unchecked except for option “C”, “Grid data already exists for image(s)” and option “D”, “Cell analysis”. Option “B”, “process a single image” can be both checked or unchecked, depending on if the user desires to process a single image or an entire folder respectively. A set of .csv files with the cell data will be created for each image

**Fig. 10.**
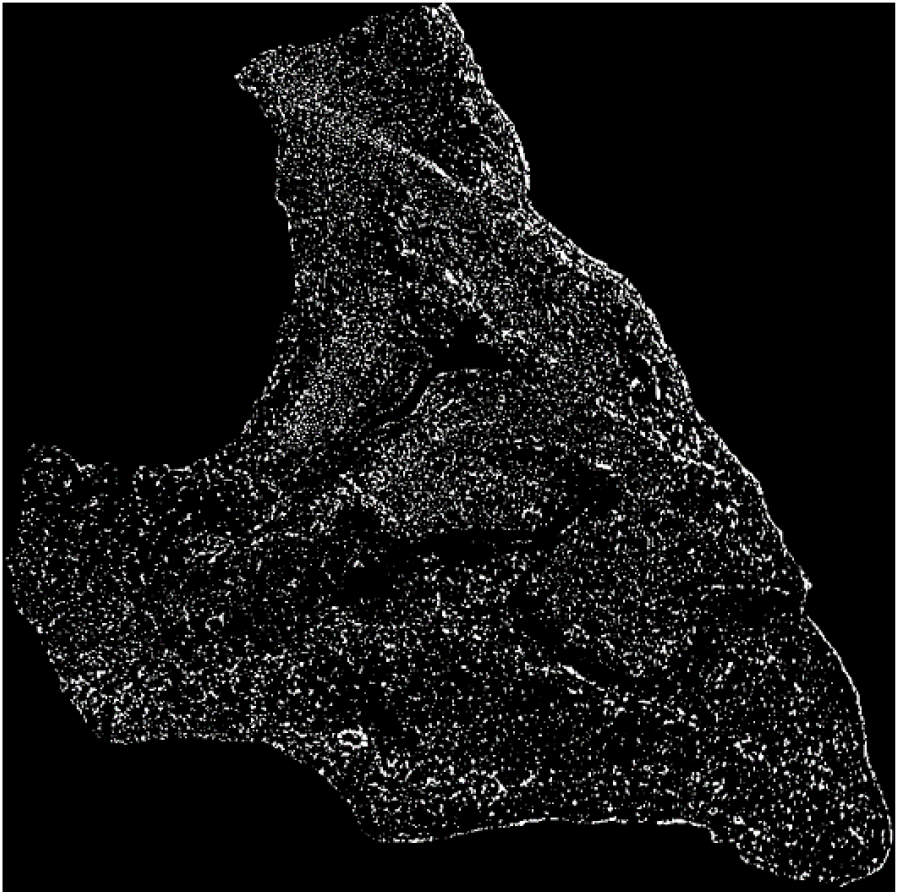
Representative thresholded image created by FinCH during cell morphology analysis.

## Data files and data file processing

### 1. FinCH outputs three types of data files

#### 1.1 Data.csv

This file contains histogram data for processed images with headers that describe the data in each column. Black pixel counts represent AOI tissue pixels that do not contain the stain representing the transcription factor for that image (negative pixels). Black pixels do not include background pixels or pixels outside of grid/image. White pixel counts represent AOI tissue pixels that contain the stain representing the transcription factor for that image (positive pixels).

There are three main sections of data file. The first column contains image name, the next three columns contain overall histogram data for the images (Total pixels | black pixels | white pixels), and the subsequent columns contain histograms of data broken up into the individual grid boxes: [[box coordinates (‘A1’), [histogram for that box as black pixels, white pixels, total pixels]], [‘B1’ [black pixels, white pixels, total pixels]], [‘C1’ …. The first column header (“Image Name”) also contains a time stamp of data collection.

#### 1.2 Grid.csv

This file contains grid information for reapplying grids to other images. It contains pixel coordinates for grids matched to corresponding images. Saving grid.csv file with AOI data enables repeated application of the same grid to the matched images. For example, when the image needs to be analysed for new histological data or in the new anatomical structure, that can be matched with the previously collected TF expression or cell morphology data in surrounding tissues of the same image.

#### 1.3 grid_*Image_name*.zip

This file is saved in the “Grids” subfolder inside the “Processed” folder and contains the coordinates of grid boxes and the grid cell ID for its corresponding image.

### 2. Cell morphology module of FInCH generates one type of data files

#### 2.1. Grid_ID.csv

This file contains information about each cell measured within each grid box. Default measures (ImageJ convention) for each cell are: Area, X coordinates, Y coordinates, Perimeter, BX, BY, Width, Height, Major, Minor, Angle, Circ., Feret, RawIntDen, FeretX, FeretY, FeretAngle, MinFeret, AR, Round, and Solidity.

### 3. Managing cell datasets

3.1 For each image, merge all .csv files into a single .csv file named with the name of the image. We used scripts in https://merge-csv.com/ and left the “Keep header (index) only in first file” option unchecked.
3.2. In many cases, whole AOI datasets of cell morphology and TF expression collected by FInCH will exceed MS Excell limits for the number of rows (~1×10^6^) and columns (~16×10^3^). It is thus recommended to create separate MS Excel files for each stain. We used Power Query in Microsoft Excel to merge all files into a single excel file for each stain.

#### Error messages and important issues

1. “Incorrect file type”. FInCH works with .tif files.
2. It is important not to skip an image while drawing the horizontal reference line in a set of open AOI images in step 1.7. Restart the script if you skipped an image.
3. “Processed/grid.csv file not found”. When reapplying the grid to a previously processed image (Step 2), the program will look for a grid.csv file associated with that image. If the file does not exist or is in the wrong folder, the program will output this error. It will then give you the option to generate new grid information. At this point FInCH can either be cancelled or new grid data can be created. If you believe a grid has already been created, you should exit FInCH and verify that the grid.csv and data.csv exist for that run and that they are in the correct folder. If files do not exist, restart FInCH and process the folder again, leaving option C unchecked.
4. Note that when collecting new data or generating a new grid, if previous csv files exist for that run, they will be overwritten. So that if you try to remeasure just one or few AOI images in the set, it is recommended that you move these files into a separate folder before the analyses.

## CRediT author statement

**Cody A. Lee, Carmen Sánchez Moreno, Alexander V. Badyaev\***: Software, Validation, Visualization, Data Curation, Writing-original draft preparation, Writing –Reviewing and Revision

## Acknowledgments

We thank Maxwell Gleason, Georgy Semenov, Sarah Britton, Renée Duckworth, Fatima Bravo, Kaitlyn Gahl, Kathryn Chenard, Chris Seliga, Robert Hollingsworth, Jakob Abtahi, Omar Puebla, and Ali Shaikh for help with collection and preparation of samples, cryosectioning, histological and molecular assays and computational work. This work was supported by the grants from the David and Lucile Packard Foundation and National Science Foundation (IBN-0218313 and DEB-1754465) to AVB and the Arthur L. and Lee G. Herbst Endowment for Innovation and the Science Dean’s Innovation and Education Fund to CSM.

## Declaration of interests

☒ The authors declare that they have no known competing financial interests or personal relationships that could have appeared to influence the work reported in this paper.

## Supplementary material *and/or* additional information

### Supplementary Note 1: Immunohistochemistry

Sections were blocked with an endogenous peroxidase (2% H_2_O_2_) for 20 minutes, washed in tris-buffered saline (TBS), blocked with 2% bovine serum albumin (BSA) plus 5% normal goat serum (NGS) in TBS for 1 hour. Avidin-Biotin blocking kit (Abcam) was used per manufacturer’s instructions before applying antibodies. Samples were incubated for 18 hrs with a primary antibody at 4°C, washed in TBS, incubated at room temperature for 1 hour with a secondary antibody, and washed again with TBS. Appropriate dilution of the primary antibody was determined by running preliminary trials with serial dilutions. Primary antibodies included anti-β-catenin (610153, 1:16,000, BD Transduction Laboratories), anti-CaM (sc-137079, 1:15, Santa Cruz Biotechnology), anti-Wnt4 (ab91226, 1:800; Abcam), anti-TGFβ2 (ab36495, 1:800, Abcam), anti-Bmp4 (ab118867, 1:100, Abcam), anti-Ihh (ab184624, 1:100, Abcam), anti-Dkk3 (ab214360, 1:100, Abcam), and anti-FGF8 (89550, 1:50, Abcam). Three different secondary antibodies (Biotinylated Goat Anti-Mouse IgG, BP-9200; Biotinylated Goat Anti-Rabbit IgG, BP-9100; Biotinylated Goat Anti-Rat IgG, all 1:200, Vector Labs) were used depending on the host primary antibodies. Negative controls were incubated with TBS instead of primary antibodies. During optimization of assays, controls were also run with just the secondary antibody to verify there was no non-specific binding. Reactions were visualized with either diaminobenzidine (DAB, Elite ABC HRP Kit, PK-6100, Vector Labs) or Vector Red Alkaline Phosphatase substrate and Vectastain ABC‐AP Kit (AK-5000, Vector Labs). Hematoxylin was used to counterstain nuclei. Each slide contained four sections, and the three slides (12 sections per embryo) were run with the following grouped antibodies: i) β-catenin, FGF8, TGFβ2 and control, ii) BMP4, WNT4, IHH and control, and iii) DKK3 and control, and CAM and control (Appendix S2). Suitable upper beak midline sections with expressed TF (*n* = 5,044) were imaged and named according to embryo ID, TF, developmental stage, and IHC run to enable automated processing.

### Supplementary Note 2: AOI selection

Cell measurements and IHC expression were conducted within AOI of the upper and lower beaks. AOIs were delineated based on landmarks homologous across developmental stages and confirmed with H&E histological staining. **Older stages, upper beak**. Start where the outer edge of the upper beak meets the top of the image. Follow the top of the image until you reach a change in tissue that marks the outer edges of the eye/brain. Continue selecting downwards, following the edge of the tissue. Select the boundary between outer eye-socket tissue and upper beak tissue, follow this line down until you reach a change in tissue. Then cut straight across the tissue to the lower edge of the upper beak (inside of the mouth). Continue to trace the edge along the inner mouth and around the tip of the beak. When you reach the egg tooth, continue to follow the outer edge. Continue along the outer edge until you meet where you started. **Older stages, lower beak**. Start where the tongue meets the lower beak. Cut straight down to the lower edge of the outer beak. Follow the edge of the lower beak all the way around until you reach where you started.

**Early stages, upper beak**. Begin at the upper outside edge of the upper beak. Start at the noticeable change in angle of the upper beak, select straight inwards to the first tissue change, follow the outer tissue of the brain/eye down until you reach the closest edge of the eye to the mouth. From the lower edge of the eye to the inner edge of the mouth you should expect quite a bit of variation. Select from the lowest part of the eye to the inner edge of the mouth just past the palatine process and then select across so that the palatine process is not included. Once you reach the inner edge of the mouth, finish selecting the upper beak.

## Notes

https://youtu.be/F-kOP2SP72E

### Competing Interest Statement

The authors have declared no competing interest.

## References

[1] A.S. Wilkins, The Evolution of Developmental Pathways, Sinauer Associates, Sunderland, MA, 2001.

[2] A.E. Kalyuzhny, Immunohistochemistry: Essential Elements and Beyond, Springer 2016.

[3] A.V. Badyaev, C.A. Lee, M.J. Gleason, G.A. Semenov, S.E. Britton, R.A. Duckworth, Tuning regulatory signaling network to cell jamming transitions can delineate population divergence in morphogenesis, BMC Biology (2024).

[4] A.V. Badyaev, The Beak of the Other Finch: Coevolution of genetic covariance structure and developmental modularity during adaptive evolution, Philosophical Transactions of the Royal Society, Biological Sciences 365 (2010) 1111–1126.

[5] A.V. Badyaev, Evolutionary significance of phenotypic accommodation in novel environments: An empirical test of the Baldwin effect, Philosophical Transactions of the Royal Society 364 (2009) 1125–1141.

[6] A. Abzhanov, W.P. Kuo, C. Hartmann, B.R. Grant, P.R. Grant, C.J. Tabin, The calmodulin pathway and evolution of elongated beak morphology in Darwin’s finches, Nature 442(7102) (2006) 563.

[7] P. Wu, T.-X. Jiang, S. Suksaweang, R.B. Widelitz, C.-M. Chuong, Molecular shaping of the beak, Science 305 (2004) 1465–1466.

[8] R. Mallarino, O. Campàs, J.A. Fritz, K.J. Burns, O.G. Weeks, M.I. Brenner, A. Abzhanov, Closely related bird species demonstrate flexibility between beak morphology and underlying developmental programs, Proceedings of the National Academy of Sciences of the United States of America 109 (2012) 16222–16227.

[9] P. Wu, Ting-Xin Jiang, Jen-Yee Shen, Randall Bruce Widelitz, C.-M. Chuong, Morphoregulation of Avian Beaks: Comparative Mapping of Growth Zone Activities and Morphological Evolution, Developmental Dynamics 235 (2006) 1400–1412.

[10] A.L. Potticary, E.S. Morrison, A.V. Badyaev, Turning induced plasticity into refined adaptations during range expansion, Nature Communications 11 (2020) 254.

[11] J. Schindelin, I. Arganda-Carreras, E. Frise, V. Kaynig, M. Longair, T. Pietzsch, S. Preibisch, C. Rueden, S. Saalfeld, B. Schmid, Fiji: an open-source platform for biological-image analysis, Nature methods 9(7) (2012) 676–682.

[12] A.C. Ruifrok, D.A. Johnston, Quantification of histological staining by color deconvolution, Anal Quant Cytol Histol 23 (2001) 291–299.

